# Transient Aurein 1.2 pores in prokaryotic model bilayers explored by coarse-grain molecular dynamics simulations: as glycolipid content increases, pore lifetime decreases

**DOI:** 10.1101/2023.01.24.525384

**Authors:** GE Balatti, MF Martini, M Pickholz

## Abstract

The Aurein 1.2 is an anuran antimicrobial peptide (AMP) with a proven lytical activity against bacterial membranes. Previously, we found a differential action of aurein by both experimental and computational methods. This differential action was over membranes of two related probiotic strains, where the main differences between membranes were the number of glycolipids on lipid composition. In the present work, we focused on the differential behavior of the aurein 1.2 at its interaction with bacterium model membranes with different amounts of glycolipids in their composition. We carried out extensive molecular dynamics (MD) simulations by using the coarse-grain force field MARTINI and raising up differential mixtures of phosphatidylglycerol (PG), phosphatidylethanolamine (PE) and monogalactosylglycerol (MG). We found a correlation between the presence of MG in PG/PE mixtures and the difficulty of aurein to stabilize pore structures, suggesting an AMP-resistance factor encoded in the lipid composition of the membrane. Through this study, we hope to shed light on a possible resistance mechanism to AMPs related with the glycolipid content of diverse types of bacterial membranes.

## 1 INTRODUCTION

Antimicrobial peptides (AMPs) are present in virtually all domains of life, exerting a broad-spectrum action, and positioning as molecules with a high-potential pharmaceutical application (Zhang *et al*., 2019; Moretta *et al*., 2022; Wang *et al*., 2022). In effect, different AMPs are being investigated as new families of antibiotics, antivirals (Elnagdy and AlKhazindar, 2020; Mousavi Maleki, Rostamian and Madanchi, 2021), anticancer (Zhong *et al*., 2021; Valenti *et al*., 2022), antibiofilm (Hancock, Alford and Haney, 2021; Lopes *et al*., 2022) and as immunomodulators (Martell *et al*., 2021), among others. The broad-spectrum, and the presence into diverse life forms made the AMPs active molecules with remarkable structural and functional diversity. Even though some AMPs can inhibit biological processes, or interact with intracellular targets (Haney, Straus and Hancock, 2019; Raheem and Straus, 2019), most AMPs affect the lipid bilayers, disrupting the physical integrity of bacterial cell membranes at different degrees and increasing the membrane water permeability (Huang, Huang and Chen, 2010; de la Fuente-Núñez *et al*., 2016; Li *et al*., 2021; Zhang *et al*., 2021). In terms of lytical activity, two main general mechanisms are described for AMPs: the “carpet-like” mechanism, where AMPs disrupt the membranes leading to micellization; and the pore formation, where membrane permeability increases preserving the overall membrane integrity. These pores encompass two principal types of structures: the “barrel-stave” structure, with the lipid headgroups remaining in the same plane; and the “toroidal” structure, where the headgroups of the lipids are forming an arc, connecting both leaflets of the membrane along the pore structure (Li *et al*., 2021).

From a structural standpoint, the cationic α-helical peptides make up an important and well-studied subgroup of AMPs. These AMPs adopt the secondary structure after interaction with lipid bilayer, being unstructured in an aqueous solution. The amphipathicity and the net positive electrostatic charge are key features of these molecules for the interaction with lipid bilayers, and the subsequent membrane properties alterations in terms of permeability, curvature, or thickness (Huang, Huang and Chen, 2010; Hollmann *et al*., 2016; Stone *et al*., 2019).

The aurein 1.2 (GLFDIIKKIAESF-NH_2_) belongs to this subgroup of AMPs and can be found on the skin of the *Litoria raniformis* and *L. aurea* Australian Tree Frogs, like other related peptides as maculatin and citropin. The physico-chemical properties of the aurein are compatible with the AMPs, including net charge (+1), average hydrophilicity (0.0 using the Hopp and Wood’s scale (Hopp and Woods, 1981)), percentage of hydrophilic (38%) and hydrophobic moment (μH=6.77). The helical wheel projection (Balatti *et al*., 2017) exhibits the polar and non-polar surfaces well delimited and equivalent in terms of extension, typical of amphipathic peptides. The secondary structure was studied by Circular Dichroism (CD) showing either unstructured or helical conformation depending on the aqueous or lipid environment (Fernandez *et al*., 2012).

The molecular mechanism and the preferential targets of aurein have not been fully elucidated. In previous works, we unveiled details on the molecular mechanism of action of this peptide on lipid membranes by MD simulations. Moreover, the simulations of aurein with POPC membranes exhibited a well-defined amphipathic behavior with a structuring compatible with pore formation and a characterized helical orientation (Balatti *et al*., 2017). In addition, we observed a similar behavior with bacterial gram-positive model membranes composed by 3:1 POPG:POPE mixtures (Balatti, Martini and Pickholz, 2018). Furthermore, we reported a differential action of aurein against two related probiotic strains with different lipid composition: *Lactobacillus delbrueckii sub. Lactis* (CIDCA133) and *L. d. subsp. Bulgaricus* (CIDCA331). The strain CIDCA133 showed a higher minimal inhibitory concentration (MIC) and cell viability (flow cytometry experiments) compared to the strain with CIDCA331. In addition, CIDCA331 was significantly inhibited by aurein in cell culture kinetics studies in comparison with CIDCA133 (Szymanowski *et al*., 2019). These results were also extended to liposomes derived from lipid extracts of these two strains, where the liposomes with high amount of glycolipids were refractory to the action of aurein in fluorescence release experiments (Szymanowski *et al*., 2019). The resistance showed by this strain was also studied by MD simulations, where the stability of pre-formed pore structures was evaluated, with equivalent results (Balatti *et al*., 2020).

In this regard, the key difference between membrane compositions of both strains is the glycolipid percentage. While the glycolipid/phospholipid ratio (GL/PL) for CIDCA331 is 3.08, the GL/PL for CIDCA133 is 23.33, exhibiting a remarkable number of saccharides in their lipid bilayer. Since CIDCA133 was refractory to aurein action, the potential passive-resistance effect associated with the presence of glycolipids groups in the membrane composition of a gram-positive bacterial bilayer model is analyzed here. In this way, we considered mixtures of phosphatidylglycerol (PG), phosphatidylethanolamine (PE) and monogalactosylglycerol (MG) lipids with increasing concentrations of the glycolipid content. The aim of the work is to compare the action of aurein over the different model membranes and with our previous work on PG:PE lipid structures in the interaction with the same peptide.

## 2 METHODOLOGY

### 2.1 Molecular dynamics systems set-up

The MD simulations were performed with GROMACS 2018.3 (Van Der Spoel *et al*., 2005; Hess B Kutzner C van der Spoel D, 2008; Abraham *et al*., 2015) software package; while CG parameters were taken from the MARTINI force field for phospholipids (Marrink, de Vries and Mark, 2004; Marrink *et al*., 2007), glycolipids (López *et al*., 2013), aminoacids (De Jong *et al*., 2013) and the polarizable water model (PW) (Yesylevskyy *et al*., 2010). The coordinates of the peptide were obtained from the NMR structure (code 1VM5 (Wang, Li and Li, 2005)) present in the Protein Data Bank (Berman *et al*., 2000). To generate the peptide topology, we used the *martinize* script, version 2.6. The peptide coordinates were also used to determine the secondary structure assignment by using the *Define Secondary Structure of Proteins* (DSSP) program (Kabsch and Sander, 1983; Joosten *et al*., 2011). We chose to model the peptide with an amidated terminal, by simply switching the residue net charge from - 1.0 to 0.0, since previous works demonstrated the relevance of the amidated C-terminal in the aurein 1.2 for the function (Shahmiri, Enciso and Mechler, 2015; Shahmiri and Mechler, 2020).

We built up 5 different model membranes of bacteria based on 1-palmitoyl-2-oleoylphosphatidylglycerol (POPG), 1-palmitoyl-2-oleoyl-phosphatidyl-ethanolamine (POPE), and 1-oleoyl-2-palmitoyl-3-alpha-galactosyl-glycerol (OPMG) mixtures. Each model membrane differed on the OPMG glycolipid percentage over the total lipids, while the POPG:POPE relation remained constant to emulate a generic gram-positive model membrane, as shown in Table 1. To simplify notation, we call each system with the glycolipid percentage (i.e., OPMG75 corresponds to the system with 75% OPMG). In particular, simulations of a previously reported **OPMG00** (0% OPMG) system were extended up to 20 microseconds. The systems were studied in the presence of 32 molecules of aurein peptides.

**Table 1.**
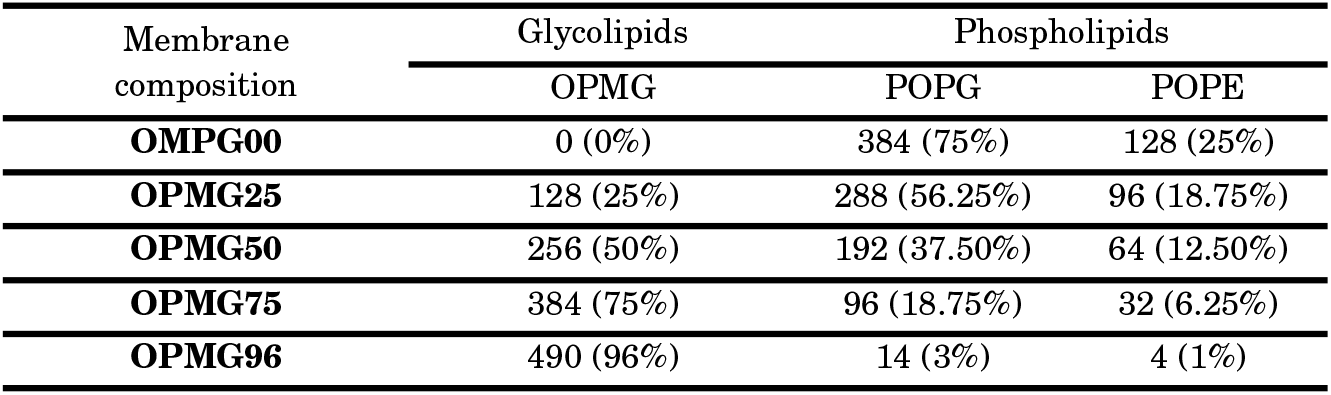
Membrane compositions

The membranes were built using the *insane* script (Wassenaar *et al*., 2015) and carefully equilibrated by increasing gradually the temperature from 10 K to 310 K. The peptides were placed inside the hydrophobic core of the different bilayers and equilibrated again by slowly increasing of temperature and the time-step to the production-run conditions. Finally, MD production simulations were performed with a time-step of 20.0 fs and both Coulomb and Van der Waals distances cut-offs were set at 1.2 nm, in a NP_XY_P_Z_T *ensemble*, using periodic boundary conditions (PBC). The temperature was controlled at 310 K with the V-rescale thermostat (Bussi, Donadio and Parrinello, 2007) at a time constant of 1.0 ps. The pressure was coupled by using the semi-isotropic Parrinello-Rahman barostat (Parrinello and Rahman, 1998) (xy and z pressures were coupled independently at 1 bar) using a compressibility of 4.5×10^-5^ bar^-1^ with a time constant of 1.0 ps in a rectangular simulation box.

### 2.2 Trajectory analysis

For the snapshots, the molecular visualization program VMD (Humphrey, Dalke and Schulten, 1996) was used. Trajectory plots and electron density profiles (EDPs) were calculated with the *gmx traj* and *gmx density* tools from the GROMACS suite; using the number of electrons for each coarse grain bead for electron density calculation.

To calculate the presence of water inside the membrane, we estimated the amount of water at the hydrophobic core, by the integration of the electron density water profile between −1 and +1 of the Z axis, and we correlate the integration value of the total water density, according to the next equation:

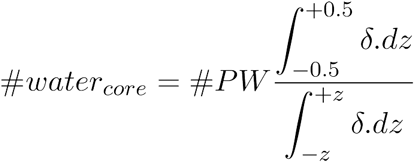

where #PW is the total beads of water molecules, *δ* is the electron density of water molecules and *z* the axis coordinate along the bilayer.

To estimate the pore lifetime, we followed a similar strategy, but in a narrow zone of hydrophobic core, integrating between z=-0.5 and z=0.5 and then correlating with the integration value of the total peptide density, according to the next equation,

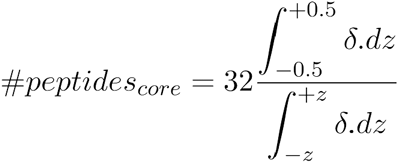

where 32 is the total number of peptides, *δ* is the electron density of the peptides and *z* the axis coordinate along the bilayer. When #*peptides_core_* < 0.5 we considered that the pore no longer exists.

In addition, to calculate peptide molecules inside the membrane we defined a “hydrophobic region” of 1 nm from each side of the bilayer center (resulting in an area with height = 2 nm) and counting the peptide BB beads with the *select* tool of the suite.

And finally, the Z-axis coordinate time evolution for the peptides and the peptide terminals were calculated using the *gmx traj* tool and defining the bilayer limits with the Phosphate (PO4) and Galactose (C1, C2, C3) beads considered as “lipid headgroups”.

*Grace* (XMGrace, 2018) and the *matplotlib* library of Phyton (Hunter, 2007) were used to plot all the data obtained.

## 3 RESULTS

The formation of pore structures is widely postulated in literature as the main mechanism of action of AMPs in membranes. Despite the fact MD simulations exhibited results in agreement with experimental results, it is challenging to get the overall process of pore formation through MD simulations starting from peptides in solution, especially with aurein. This is due to two main factors: the collective nature of pore formation and the long timescales enveloped in the process (Rzepiela *et al*., 2010; Miyazaki, Okazaki and Shinoda, 2019). Nevertheless, MD simulations showed reliable results in characterizing transient pores in lipid bilayers (Chen *et al*., 2019). In particular, our previous works overcame these limitations starting from initial configurations where the peptides were originally placed inside the membrane.

Using these initial conditions, we accessed aurein pore formation in POPC (Balatti *et al*., 2017) and POPG/POPE (Balatti, Martini and Pickholz, 2018) membranes using CG models. In addition, and taking these results into account, we explored the stability of aurein pores in more complex, atomic-detail membranes containing glycolipids. In this last case, we noticed that the pore stability was related to the presence of glycolipids in the bilayer, driven mainly by protein-lipid hydrogen bond and electrostatic interactions. These findings led us to question the dependence of pore stability on the glycolipid molar percentage in lipid mixes. So, here we explored the aurein 1.2 aggregation inside bilayers with different lipid compositions using a coarse grain model. The bilayer compositions include 3 lipid types: POPG, POPE and OPMG. As we already mentioned, the different bilayers were named as *OPMGNN*, where *NN* is the mole percent of glycolipid in the mix, as shown in Table 1. In all studied cases, the aurein 1.2 peptides were initially placed in the hydrophobic region of the bilayer. Here, transient pore formation was observed in all glycolipids containing bilayers. To investigate the pore lifetime for each case, the simulations were run until the pore was disassembled. A particular case was the OPMG00 (without glycolipids) bilayer. In this case, we extended the simulation beyond 20 μs and the pore remained stable.

We illustrate the behavior of the systems containing OPMG lipids using the OPMG75 system as a representative case. Figure 1 shows representative snapshots of this system over time. Initially, peptides are placed inside the bilayer (Fig. 1A). From this initial condition, peptides aggregated inside the bilayer in a few nanoseconds. This fact is illustrated at 200 ns (Fig. 1B). Around 500 ns, migration of single peptides from the aggregate to the lipid interphase is observed (Fig. 1C). Fig. 1D shows that at this time all peptides are found at the interphase, where they remain at the end of the simulation (2 μs for this case, Fig 1E).

**Figure 1.**
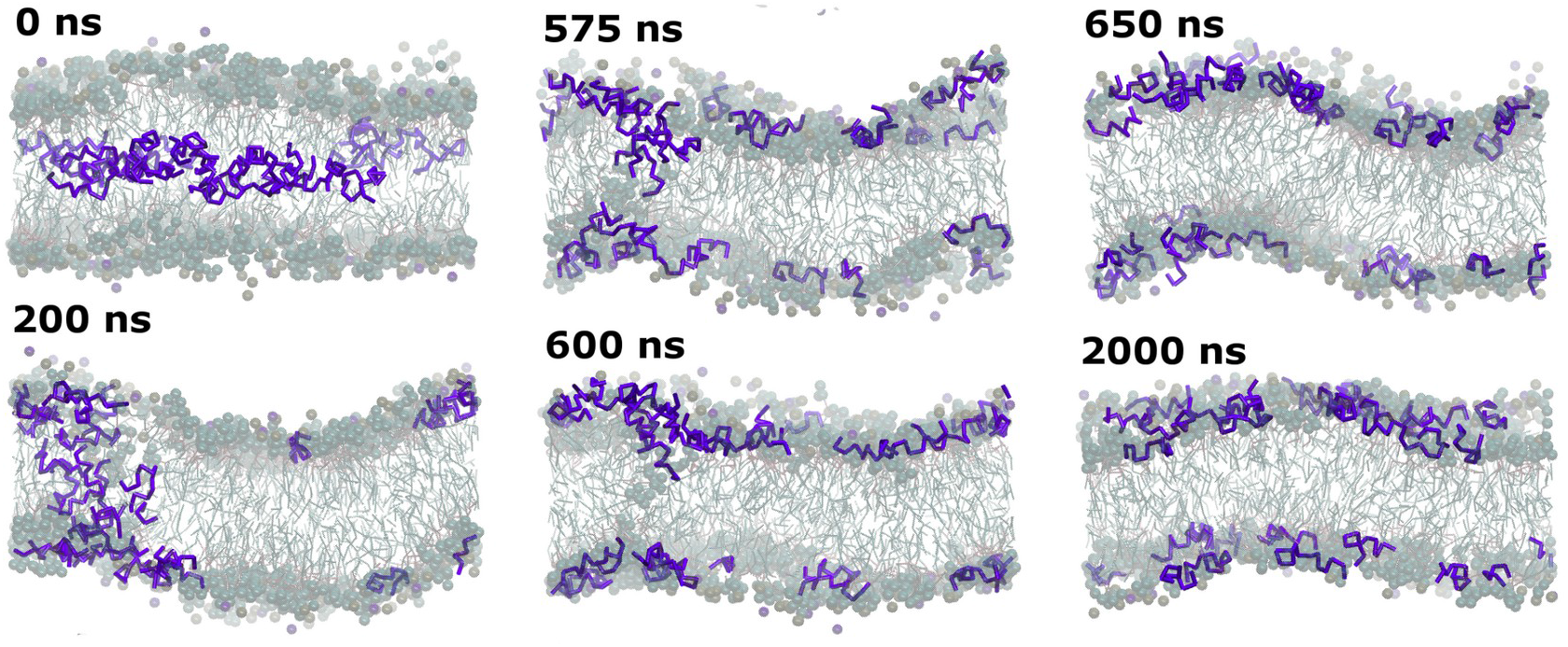
Representative snapshots that show the progression of the pore formation and destabilization in the OPMG75 membrane system over the simulation time. Peptide backbones are depicted in blue, with the polar lipid headgroups represented by balls.

Next, we compared some characteristics of the different systems that allow us to evaluate the evolution of the formation and stability of the pores depending on the number of glycolipids. Figure 2 shows the first 2 μs of z-axis trajectories of the center of mass of each of the 32 peptides (in green) of all the systems (from top to bottom OPMG96, OPMG75, OPMG50, OPMG25 and OPMG00 systems). As a reference, we have included in these figures the average position of the bead that represents lipidic PO_4_^2-^ headgroups (in black). The peptides in the membranes with high content of glycolipids quickly migrated to lipid interphases (Fig. 2 A, B y C). On the other hand, the peptide aggregates remain stable, on this timescale, in membranes with low glycolipid content (Fig. 2 D and E).

**Figure 2:**
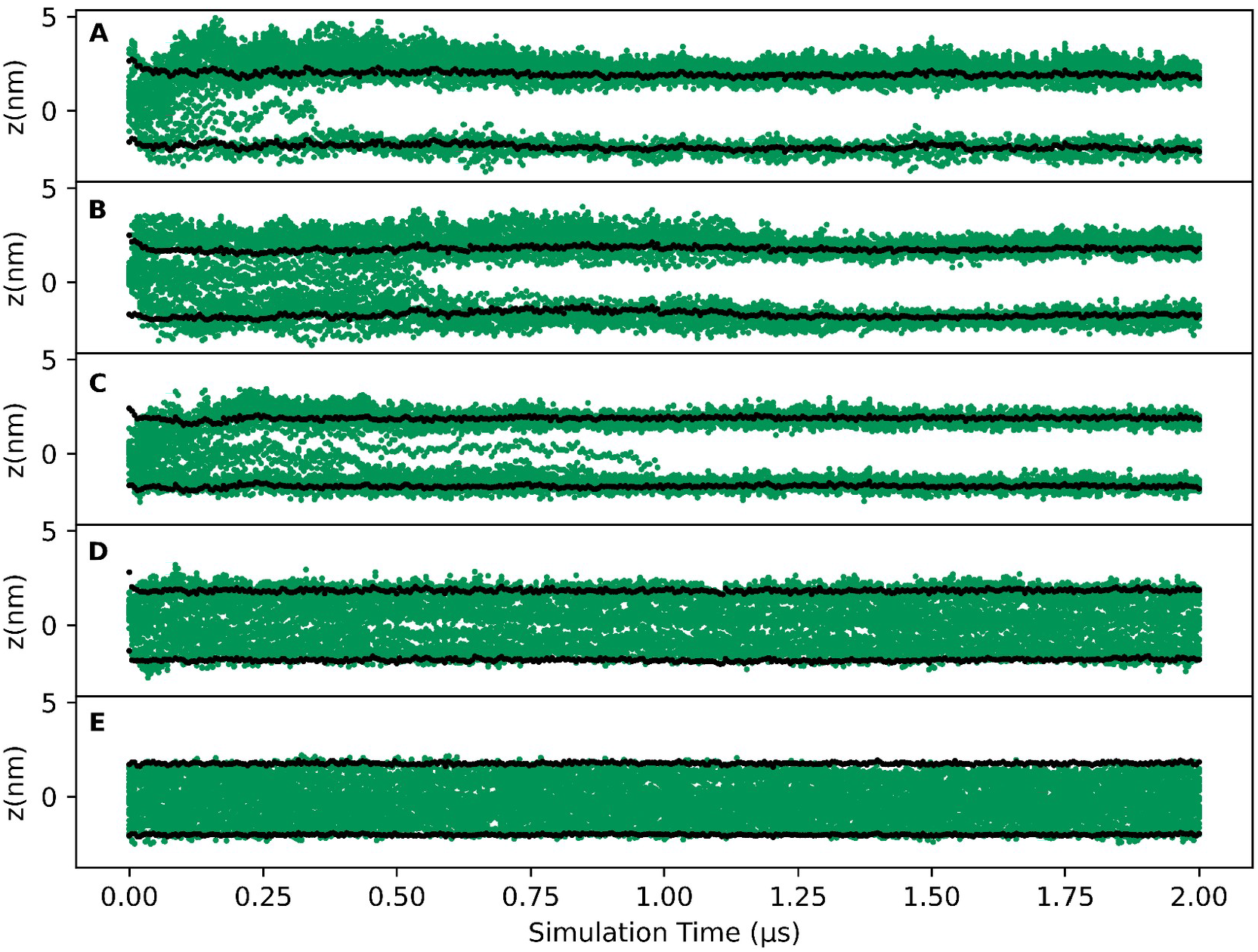
Z-axis trajectories of the center of mass of each of the 32 peptides (in green) for the OPMG96, OPMG75, OPMG50, OPMG25 and OPMG00 cases (**A** to **E**, respectively). Black lines represent the average z coordinate positions of the phosphate group beads of each monolayer, as a membrane reference

This differential peptide behavior was quantified by calculating the number of peptides inside the bilayer. Figure 3A shows the number of peptides inside the bilayer plotted as a function of time for all the systems, (during the first 2 μs of the simulation run). Using this criterion, we can estimate the lifetimes of peptide aggregate structures within the membranes. For instance, the lifetime for the peptide aggregates in the OPMG96 systems was noticeably short (~0.3 μs) and it raised progressively for the membranes with a smaller number of glycolipids (~0.5 − 1 μs for OPMG75 and ~1 μs for OPMG50). For the OPMG25 and OPMG00 systems (low glycolipid molar fraction), the peptide aggregates were stable, at least in the first 2 microseconds. To study the long-term stability of pores in these membranes beyond the 2 μs, we extended the simulations of these systems. For the OPMG25 case, the pore lifetime was ~6 μs. On the other hand, the pore remained stable for at least 22 μs for the systems without glycolipids. The lifetimes of the aggregates inside the bilayer are summarized in Table 2. A remarkable finding was that this relationship between glycolipid content and pore lifetime remains when the peptide concentration was reduced by half. Nevertheless, the overall lifetime also decreased (data not shown).

**Figure 3:**
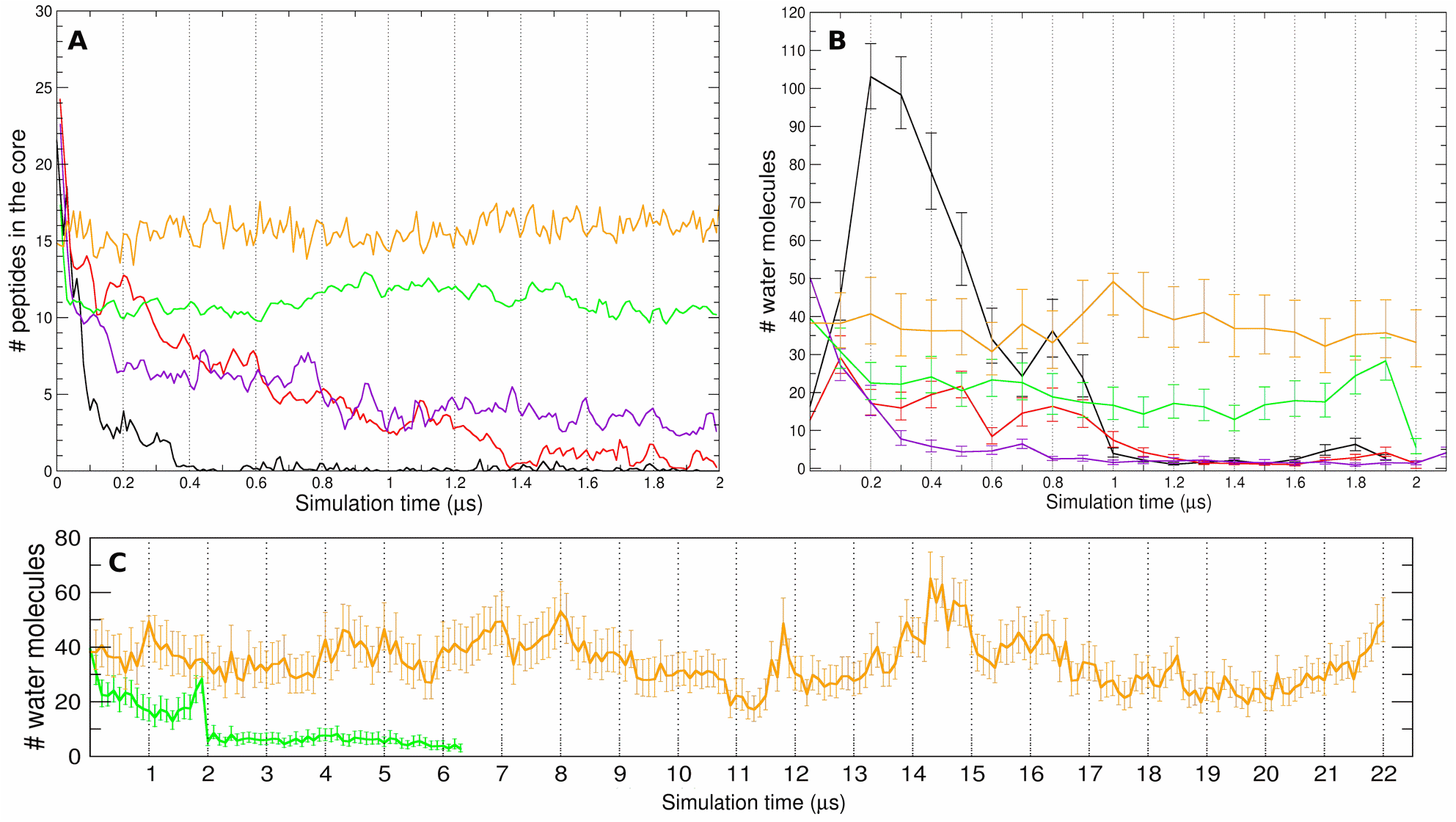
Presence of aurein molecules (**A**) and water molecules (**B**) in the hydrophobic core of the bilayer as a function of time for OPMG96 (black) OPMG75 (red), OPMG50 (purple), OPMG25 (green) and OPMG00 (orange) cases. Number of water molecules (**C**) in the hydrophobic zone of the bilayer over the extended time of 22 μs for OPMG00 (orange) and 6.5 μs for OPMG25 (green).

**Table 2.** Detailed information of simulated systems with pore structures lifetimes

| Membrane composition | System configuration | Run Time (with replica) | Permanence Time |
| --- | --- | --- | --- |
| **OPMG00** | 512 lipids + 32 aureins + | 24.00 μs | 24 μs |
| | 13119 PW + 384 NA + 32 CL | 22.00 μs | 22 μs |
| **OPMG25** | 512 lipids + 32 aureins + | 6.60 μs | 6.15-6-20 μs |
| | 13439 PW + 292 NA + 36 CL | 7.00 μs | 6.0 μs |
| **OPMG50** | 512 lipids + 32 aureins + | 2.00 μs | 0.95-1.00 μs |
| | 13418 PW + 244 NA + 84 CL | 2.00 μs | 1.00-1.10 μs |
| **OPMG75** | 512 lipids + 32 aureins + | 2.00 μs | 0.50-0.55 μs |
| | 10809 PW + 95 NA + 31 CL | 2.00 μs | 1.00-1.05 μs |
| **OPMG96** | 512 lipids + 32 aureins + | 2.00 μs | 0.30-0.35 μs |
| | 10764 PW + 86 NA + 104 CL | 2.00 μs | 0.25-0.30 μs |

To further explore peptide aggregates, we looked at two characteristics: their hydration and their organization. Water molecules were found inside the aggregates, which enables them to be called pores. We quantified the number of water beads inside the pores as a function of time. To estimate the water presence, we calculated the area under the curve of the water Electron Density Profiles (EDP) through a Simpsons defined integration between a hydrophobic core region of 2 nm (Fig. 3B). As expected, the water beads inside the bilayer decrease in the systems as the aggregate disassembles. The number of water molecules remains constant during the time shown in this figure. Figure 3C shows the evolution of the water number inside the core over long-term simulations for the OMPG25 and OMPG00.

To characterize the peptide behavior as a function of time within the bilayer, the z-axis trajectories of **NT** (green) and **CT2** (blue) beads were analyzed up to 2 μs (Figure 4). As a general observation, peptides have a preferential orientation along their longitudinal axis, with NT terminals (blue points) facing the interphase and CT2 terminals closer to the hydrophobic core (red points). This effect becomes clearer for the OPMG25 and OPMG00 systems: CT2 beads remained inside the hydrophobic core and the NT beads were inside the lipid interphase. Comparable results were obtained for the orientation of the aureins involved in pores inside in POPC and POPG:POPE membranes (Balatti *et al*., 2017; Balatti, Martini and Pickholz, 2018).

**Figure 4:**
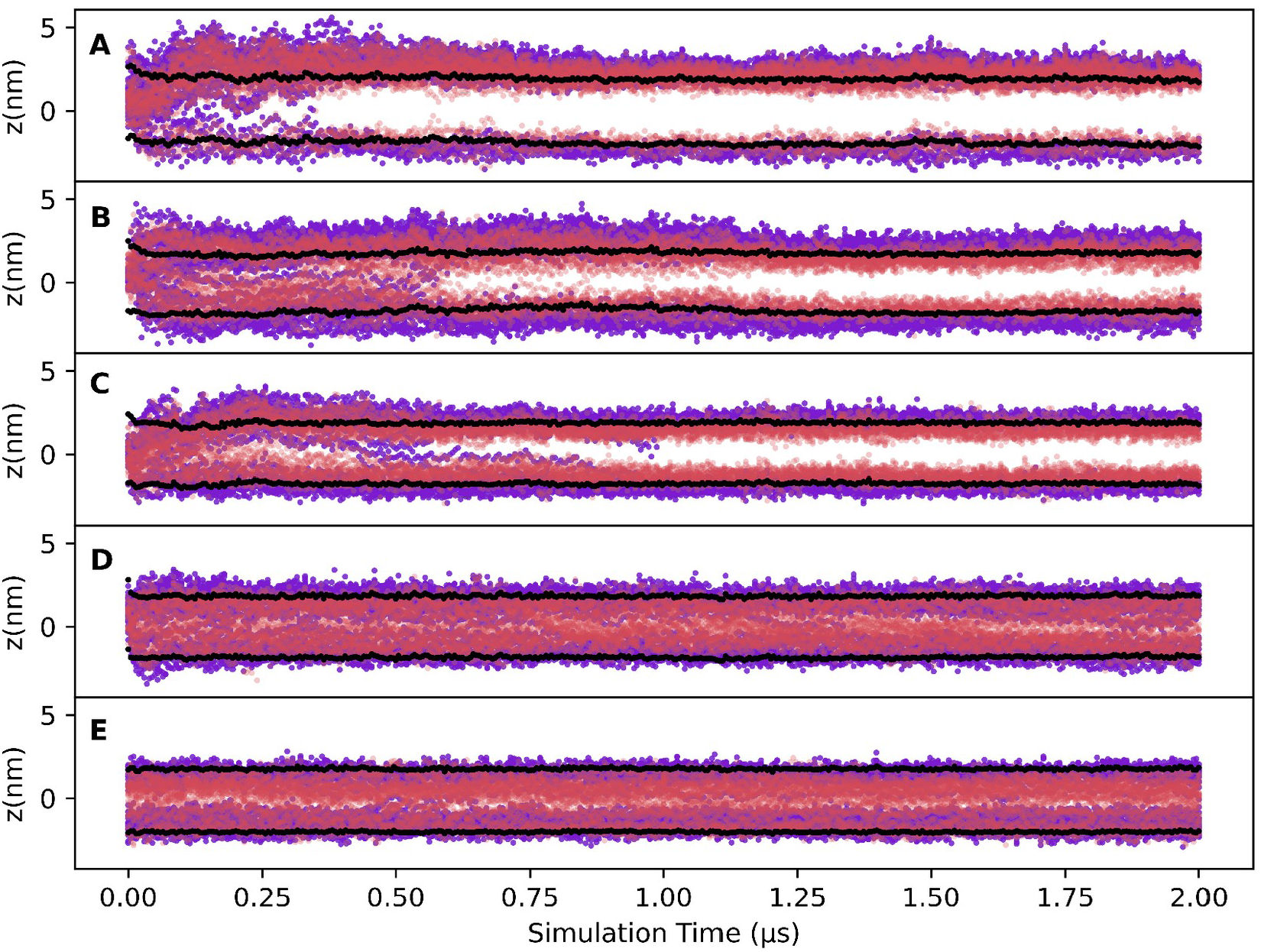
Z coordinate time evolution of amide CT2 (red) and amine NT (blue) beads as a function of time for the OPMG96, OPMG75, OPMG50, OPMG25 and OPMG00 cases (**A** to **E**, respectively). Black lines represent the z coordinate average of the phosphate groups of each monolayer, as a membrane reference

Regarding the system without glycolipids (OPMG00), we have previously reported it for one microsecond (Balatti, Martini and Pickholz, 2018). Here, we extended this system simulation to 22 μs which leads to reinforcing the results of the stability of the pore formed. Figure 5 shows a representative snapshot of the OPMG00 structure at the end of the large simulation (22 μs). This snapshot reveals the pore structure inside the lipid bilayer. From a lateral view (Fig. 5A), many water beads can be observed (represented as small blue balls) inside the hydrophobic core of the bilayer, along with some lipid headgroups (represented as big balls). A top view of this structure (Fig. 5B) shows an internal channel that allows water beads access to the core. Here, the peptides are depicted with green balls to represent the polar residues and with red ones for the non-polar residues. This representation was made with the aim to remark the great polar surface present facing the central axis of the pore, allowing water permeation through the membrane. This type of organization, with the polar surface facing the central axis and the apolar surface interacting with the lipid tails, was also observed in previous work with POPC and POPG/POPE membranes (Balatti *et al*., 2017, 2020; Balatti, Martini and Pickholz, 2018).

**Figure 5.**
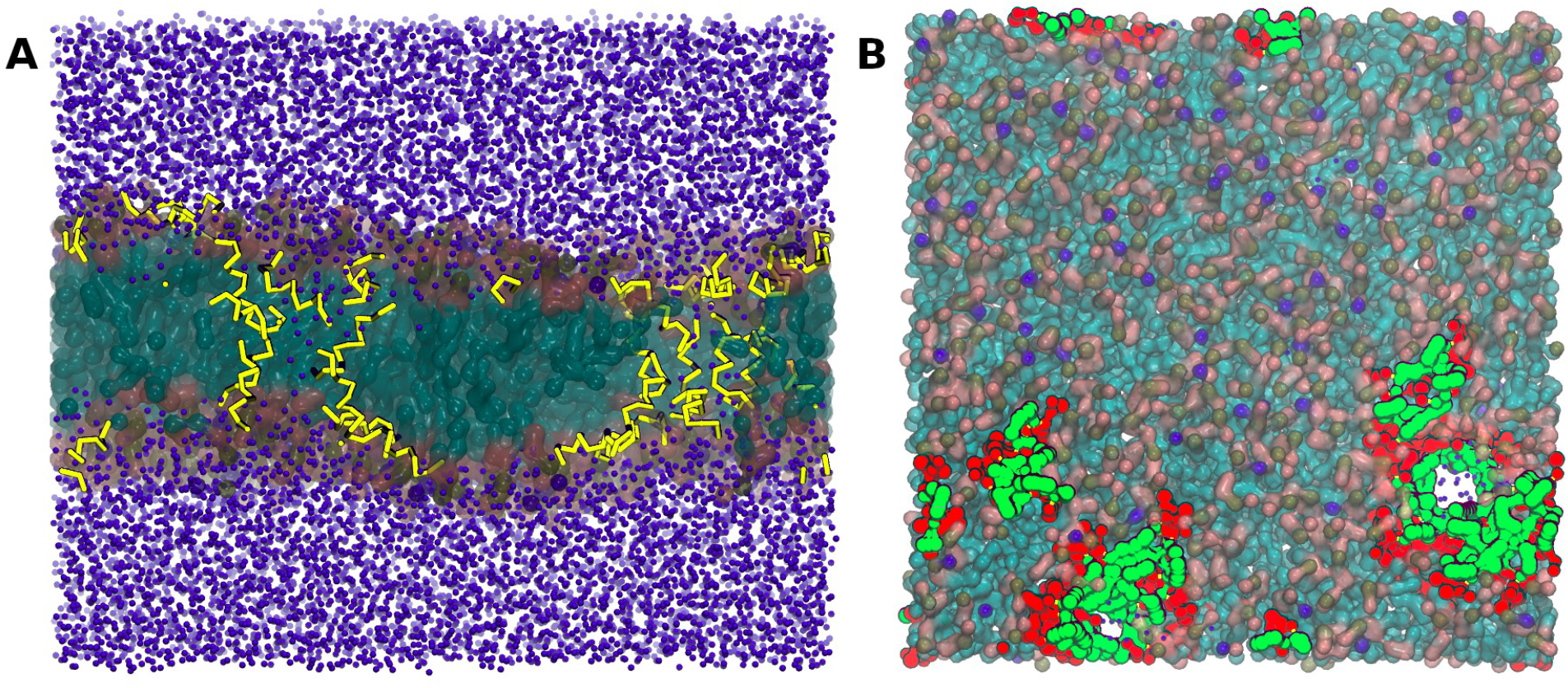
Final snapshot of the OPMG00 membrane with pore structure at 22 *μ*s from a side (**A**) and a top (**B**) view. In A: the peptide backbones are depicted in yellow, while in B: the peptides are depicted by their polar (green) or non-polar (red) residues. The water molecules are depicted only in figure A, and are represented as blue spheres.

A remarkable feature is the presence of glycolipid heads inside the bilayer when the aggregates of AMPs are present. This behavior is observed in all the systems, as could be observed from the z-axis trajectories of the glycolipid head for the OPMG75 and OPMG25 cases (Figure 6). The galactose headgroup of the glycolipids exhibited a strong tendency to migrate to the hydrophobic core during the pore residence time.

**Figura 6:**
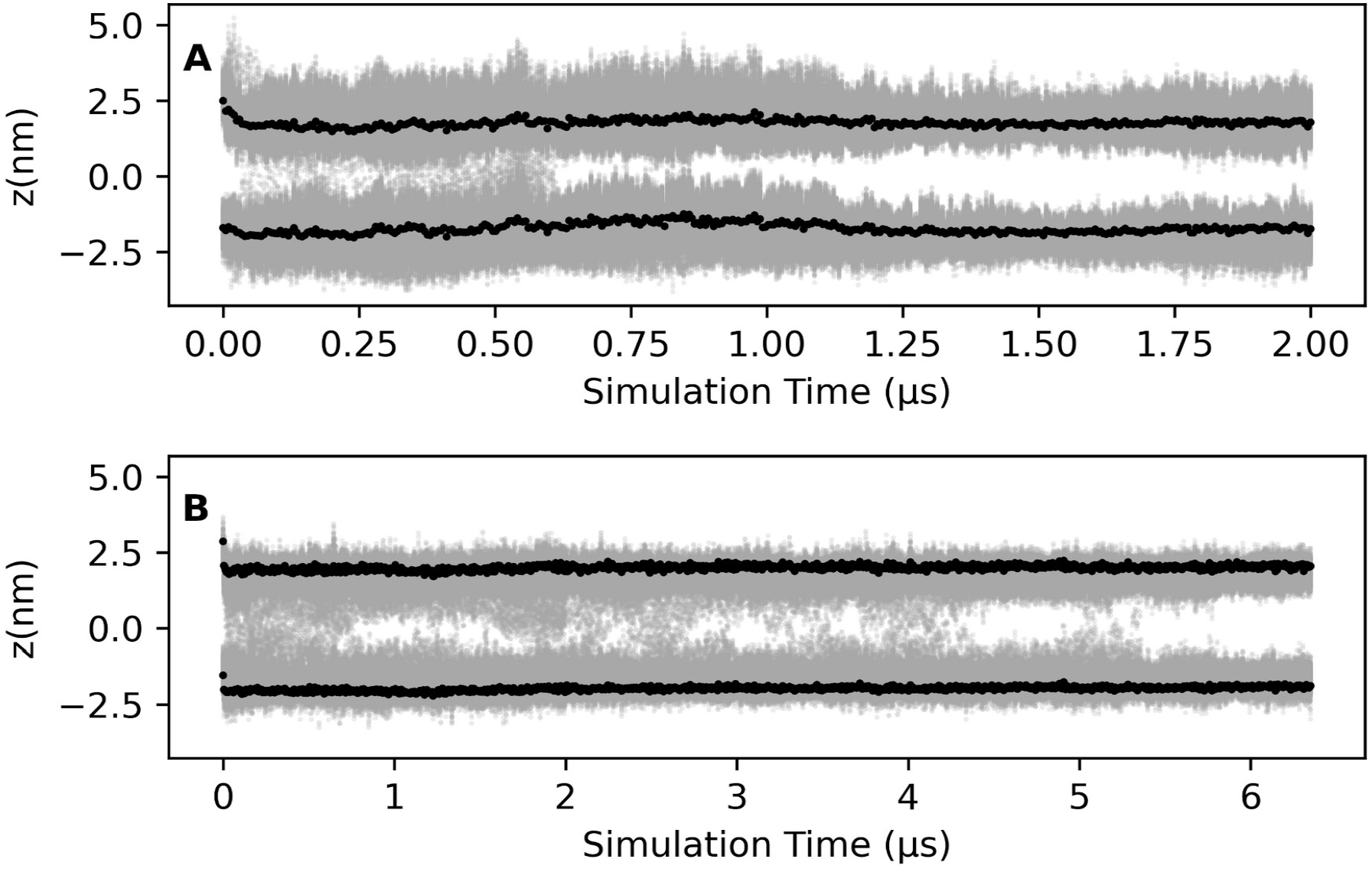
Position of galactose ring COM (gray) over the time for the OPMG75 (top) and OPMG25 (bottom) cases. Black lines represent the positions of the phosphate groups of each monolayer, as a membrane reference.

## 4 DISCUSSION

The aim of this article was to study from a molecular point of view a potential passive mechanism of bacteria to avoid the action of antimicrobial peptides, based on the presence of glycolipids on their membrane composition. The link between glycolipids presence and AMP resistance was suggested previously based on both experimental and computational observations from the probiotic strains CIDCA133 and CIDCA331 (Hugo, De Antoni and Pérez, 2010; Hugo *et al*., 2012; Szymanowski *et al*., 2019; Balatti *et al*., 2020). Here, we raised up a set of model bilayers with different amounts of glycolipids, and we ran MD simulations with the aurein AMP inside the hydrophobic core of the membranes. We observed the formation of pore structures with similar structural features to the previously simulated PC and PG/PE membranes, like a clear NT/CT orientation and a good passage of water molecules (Balatti *et al*., 2017; Balatti, Martini and Pickholz, 2018). Despite this fact, the stability of these transient pore structures exhibits a remarkable dependence on the glycolipid mole fraction. Furthermore, the pore structure in the membrane without glycolipids (**OPMG00**) is remarkably stable, even beyond the 22 μs. As the mole fraction increases, the lifetime of the pores becomes shorter. Relevant factors can be mentioned at a molecular level regarding glycolipid presence in lipid composition to bring AMP resistance advantages to the bacterial membranes. The first one is the electrostatic charge of the bilayer, relevant in the initial interaction of cationic peptides with negatively charged membranes. The presence of neutral non-zwitterionic lipids reduces the net or partial negative charges, regarding negative or zwitterionic polar head groups of the lipids, available to interact with cationic peptides (Qian and Zolnierczuk, 2022). This mechanism was proposed not only for the CIDCA133/CIDCA331 membranes (Hugo *et al*., 2012) but also for other biological systems (Winkowski, Ludescher and Montville, 1996; Hale and Hancock, 2007; Jung *et al*., 2011). Also, the presence of neutral zwitterionic lipids as PC was demonstrated to attenuate the effect of aurein and other related AMPs (Ambroggio *et al*., 2005; Fernandez *et al*., 2013). As we have previously studied, the preference of aurein for membranes with PG instead of those compose of PC (Balatti *et al*., 2017; Balatti, Martini and Pickholz, 2018) or even for the 75% glycolipid CIDCA331 instead of the 96% CIDCA133 (Szymanowski *et al*., 2019; Balatti *et al*., 2020) demonstrate from both computational and experimental approaches a preference of the aurein for negatively charged membranes.

Another remarkable characteristic of the glycolipids-based membranes is the curvature. It was suggested that the curvature of some membranes with global or local curvature can counteract the curvature-inducing action of aurein (Chen and Mark, 2011; Poger, Pöyry and Mark, 2018); and the presence of glycolipids is common in membranes with high ratios of curvature, like the MGDG of the thylakoid membrane present in chloroplasts (Webb and Green, 1991). Moreover, as we studied previously, it is an important feature of the aurein way of action, and also for the maculatin, a narrow-related peptide (Balatti *et al*., 2017; Szymanowski *et al*., 2019).

In addition, glycolipids have a high capability to form hydrogen bonds. Like all the saccharides, galactose can be a donor/acceptor of hydrogen bonds through its multiple hydroxyl groups, in comparison with the single hydroxyl group of, e.g., a PG headgroup. In fact, the ΔH and temperature of phase transition in membranes with glycolipids are higher rather than in phospholipid membranes because of the hydrogen bond network that reinforces the interaction between lipid headgroups (Koynova *et al*., 1988; Kuttenreich *et al*., 1988). This aspect was evaluated via atomistic MD simulations, where the membrane CIDCA133, with 96% of glycolipids showed a great tendency to establish hydrogen bonds not only between galactose headgroups, but also with the peptides during the disassembly of the pore (Balatti *et al*., 2020). The present results, where a wide range of glycolipid presence evaluated, give insights about a possible link between glycolipids and the resistance of certain types of membrane to the action of AMPs like aurein.

## 5 CONCLUSIONS

The results exhibited here establish a link between the presence of glycolipids in membranes and their capability to avoid the action of AMPs. The identification of the molecular determinants to provide this capability can help to guide the rational design of novel antimicrobial chemotherapies.

## ACKNOWLEDGMENTS

This work was supported by the Argentinian National Scientific and Technical Research Council (CONICET).

